# JAAG: a JSON input file Assembler for AlphaFold 3 with Glycan integration

**DOI:** 10.1101/2025.10.08.681238

**Authors:** Chin Huang, Kelley W. Moremen

**Author notes:** Supplementary Document 1: Tree structure of JAAG input and output field. Supplementary Document 2: Monosaccharide CCD csv file.

## Abstract

Standalone AlphaFold 3 (AF3) models proteins, post-translational modifications, and ligands (including glycans) from a single JSON input file. Plausible glycan stereochemistry is more consistently achieved when monosaccharides are specified as Chemical Component Dictionary (CCD) entries and connected using bondedAtomPairs (BAP) syntax. Although this approach preserves glycan stereochemistry, assembling JSON input files manually is challenging due to the diversity of monosaccharides, linkages, branching, and structural complexity. To simplify this process, we developed JAAG (JSON input file Assembler for AlphaFold 3 with Glycan integration (https://biofgreat.org/JAAG and https://github.com/chinchc/JAAG)), a web-based graphical interface that streamlines AF3 JSON file creation, reduces errors, and facilitates modeling of glycans and glycan-macromolecule interactions.

## Introduction

Glycans, in free form or covalently linked to proteins, lipids, RNA, or metabolites, are critical regulators of diverse biological processes, including cell differentiation, immune recognition, oncogenesis, among many others (Varki, A. 2017). Their structural complexity and heterogeneity, however, complicate mechanistic studies and necessitate computational approaches to complement experimental analyses. Molecular docking and molecular dynamics offer models with atom-level resolution for investigating glycan-protein interactions and conformational ensembles, but such methods scale inefficiently with system size and sampling time (Woods, R.J. 2018). Recent advances in deep learning have led to the development of novel structure prediction frameworks, such as RoseTTAFold All-Atom (Krishna, R., et al. 2024), AlphaFold 3 (AF3) (Abramson, J., et al. 2024), Chai-1 (Chai Discovery, et al. 2024), and Boltz-2 (Passaro, S., et al. 2025), that offer accelerated predictions at reduced computational cost.

In contrast to the limited ligand input options available on the AF3 server, the standalone AF3 release in late 2024 has demonstrated strong performance in modeling glycans with enzymes (e.g., glycosyltransferases, glycohydrolases, lyases) and lectins, as validated against Protein Data Bank (PDB) structures deposited after the training set cutoff date (Abramson, J., et al. 2024, Huang, C., et al. 2025). Using bondedAtomPairs (BAP) syntax and the CCD library (Westbrook, J.D., et al. 2015) to specify ligands, AF3 generates stereochemically accurate glycans. By contrast, simpler linear formats such as SMILES often yield erroneous outputs, including incorrect anomers, inverted epimer, and ring distortions (Huang, C., et al. 2025). In the hybrid BAP+CCD approach, BAP encodes inter-residue connectivity, and CCD entries specify monosaccharide stereochemistry, including anomeric status, absolute configuration, and ring conformation, thereby enabling accurate representation of glycan diversity. This strategy has been systematically evaluated in previous work (Huang, C., et al. 2025).

Despite these advantages, building JSON input files for AF3 remains complicated. For example, assembling a biantennary *N*-glycan with terminal sialylation (A2G2S2) requires selecting the appropriate CCD entries (e.g., NAG for *β*-D-GlcNAc-pyranose, BMA for *β*-D-Man-pyranose, MAN for *α*-D-Man-pyranose, GAL for *β*-D-Gal-pyranose, and SIA for *α*-Sia), identifying atom-level connection points, and defining ten BAPs for eleven monosaccharides. Scaling this to model glycoproteins such as Siglec-2 (CD22), which contains eleven *N*-glycosylation sites, requires defining over 120 BAP, while modeling dimeric interactions would demand more than 240 BAP alongside unique chain identifiers (Huang, C., et al. 2025). Such complexity renders manual assembly error-prone and inefficient for large systems.

To address these challenges, we present JAAG, a graphical interface web tool that automates JSON input generation for AF3. JAAG supports all AF3 JSON frameworks, including the most recent version 4, integrates automatic conversion of glycans from SugarDrawer (Tsuchiya, S., et al. 2021) or GlycoCT (Herget, S., et al. 2008) into BAP+CCD syntax, and provides seamless access to UniProt (2025), GlyTouCan (Tiemeyer, M., et al. 2017), and GlyGen (York, W.S., et al. 2020) through application programming interfaces (APIs) to ensure structural and annotation consistency. By automating these processes, JAAG significantly lowers the technical barrier for glycan modeling and broadens the applicability of AF3 to glycobiology.

## Results

### Layout of the JAAG Web Tool

JAAG is an interactive web-based application designed to support the complete input settings and functions provided by standalone AF3 (https://github.com/google-deepmind/alphafold3/blob/main/docs/input.md). The tool facilitates the streamlined generation of AF3-compatible JSON input files. To enhance usability and minimize potential confusion, JAAG presents only the essential input fields upon initialization, with advanced options accessible as needed (Fig. 1). A full structural tree outlining all input and output fields is provided in Supplementary Document 1.

**Fig. 1.**
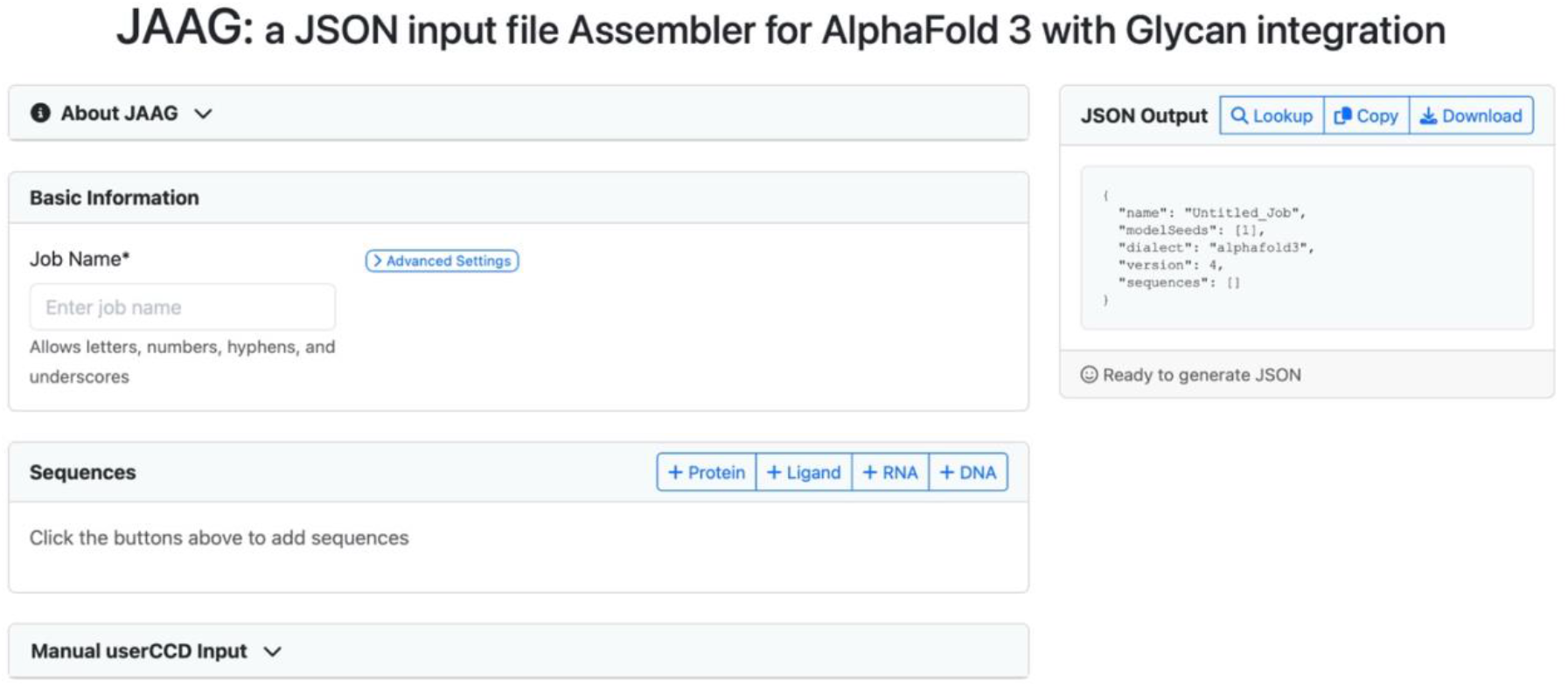
Layout of JAAG web tool at initialization. Only essential functions are displayed by default. User may add molecules of interest by selecting Protein, Ligand, RNA and DNA options within the Sequences section.

### Integrating Glycans into AlphaFold 3 JSON Input File

A curated list of CCD entries was compiled based on Symbol Nomenclature for Glycans (SNFG) (Varki, A., et al. 2015) and stereochemical properties (anomeric configuration, absolute configuration, and ring conformation) (Supplementary Document 2). GlycoCT, a widely adopted syntax, encodes stereochemical properties, linkage information, and substituents of glycans (Herget, S., et al. 2008). Leveraging this explicit representation alongside the curated CCD list enables a straightforward conversion of GlycoCT to BAP+CCD syntax.

To further facilitate glycan modeling, JAAG also incorporates SugarDrawer (Tsuchiya, S., et al. 2021), allowing users to draw glycans and convert them directly into BAP+CCD syntax. The SugarDrawer shortcut was modified to include all human monosaccharides and substituents supported in JAAG, such as sulfation, *N*-sulfation, phosphorylation, acetylation, and methylation.

JAAG also provides a “Detect Sequon” function, which identifies potential *N*-glycosylation sites within protein sequences using the N-X-S/T sequon (where X ≠ P) (Stanley, P., et al. 2022) (Fig. 2). Users can refine predictions based on experimental data by deselecting unoccupied sites. Several common *N*-glycans are available as templates, while other glycosylation types can be incorporated through a “Manual Glycosylation Site” option.

**Fig. 2.**
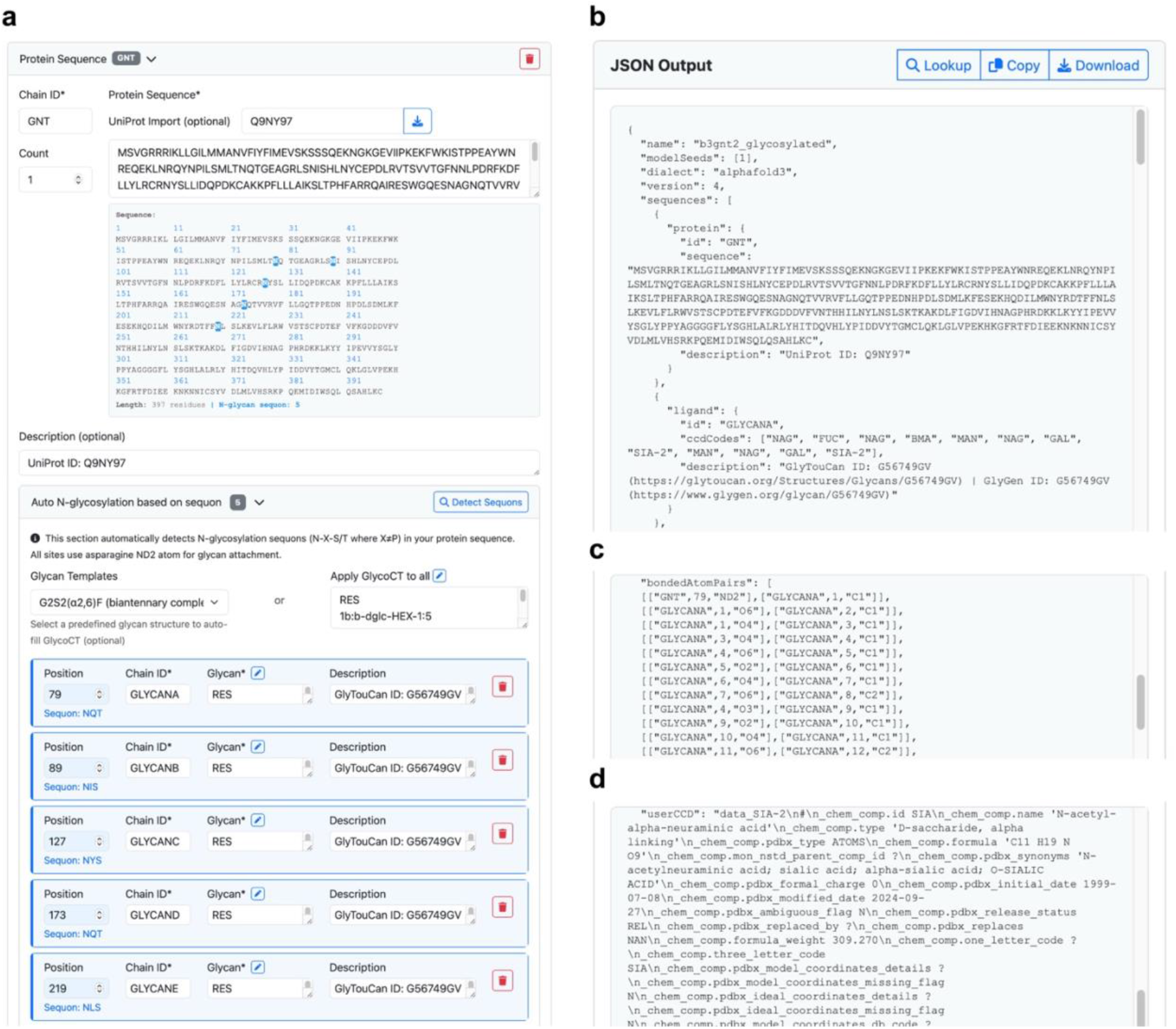
Input of fully *N*-glycosylated B3GNT2 and the corresponding JSON output by JAAG. **a**) The UniProt accession ID for human glycosylation enzyme, B3GNT2, was used to retrieve the full amino acid sequence. All *N*-glycosylation sites were annotated using the “Detect Sequons” function. A G2F glycan template was applied to all remaining sites. Only a portion of the interface is shown due to space limitations. **b**) The corresponding JSON output generated by JAAG, in which UniProt, GlyTouCan, and GlyGen identifiers were automatically embedded. Only a portion of the resulting JSON file is shown in the screenshot due to space limitations. **c**) A portion of bondedAtomPairs and **d**) userCCD for SIA generated by JAAG are shown.

### userCCD Override

In AF3, the default handling of glycosidic linkages involves the automatic removal of leaving oxygen atoms during bond formation. However, this behavior is not consistently applied to certain linkages, such as those involving sialic acids, sulfation, or phosphorylation (Huang, C., et al. 2025). JAAG addresses this by automatically triggering userCCD overrides, ensuring proper handling of Neu5Ac, Neu5Gc, sulfated glycans, and phosphorylated glycans.

### Populating Distinct Chain IDs

AF3 requires unique identifiers for each sequence defined in the JSON input. This process becomes repetitive when users model complexes containing multiple identical components, such as protein multimers, multivalent ligands, or mock lipid bilayers. To reduce redundancy, JAAG implements a “Count” function, which appends alphabetical suffixes to user-defined chain IDs and automatically propagates these identifiers to associated chains. This feature is particularly beneficial when defining glycoprotein multimers with multiple glycosylation modifications.

### API Accession Integration

To enhance reproducibility and annotation integrity, JAAG allows users to embed accession identifiers from UniProt, GlyTouCan, and GlyGen directly into the JSON input. By querying these databases, JAAG can also verify whether a drawn glycan has been previously reported, providing biological context and minimizing ambiguity during downstream analysis. This functionality additionally facilitates seamless metadata inclusion when depositing models into public repositories such as ModelArchive (Tauriello, G., et al. 2025).

## Discussion

Several barriers hinder the initiation of standalone AF3 modeling jobs, including hardware requirements (sufficient GPU and CPU capacity), successful installation of AF3 and its dependencies, permission to access DeepMind-provided model parameters, and the construction of syntactically valid JSON files that accurately encode biomolecular stereochemistry. As input complexity increases, so does the likelihood of user error.

JAAG specifically addresses the latter challenge by automating JSON assembly and reducing user-dependent variability. Its capacity to convert either GlycoCT strings or drawn glycans into a valid BAP+CCD syntax streamlines a process that is otherwise labor-intensive and error-prone. Additional features, such as sequon detection, chain ID management, and integration with glycoinformatics databases, further enhance usability, reproducibility, and data consistency. Collectively, these functionalities establish JAAG as a practical resource for the structural biology and glycobiology communities, enabling more efficient modeling of glycans using AF3.

## Acknowledgements

We thank Jordan Utley (GACRC, University of Georgia) for the maintenance of AlphaFold 3, René Ranzinger (CCRC, University of Georgia) for his valuable suggestions on JAAG, and Sujeet Kulkarni (CCRC, University of Georgia) for guidance on the GlyGen API.

## Reference

2025. UniProt: the Universal Protein Knowledgebase in 2025. Nucleic Acids Res, 53:D609–d617.

Abramson J, Adler J, Dunger J, Evans R, Green T, Pritzel A, Ronneberger O, Willmore L, Ballard AJ, Bambrick J, et al. 2024. Accurate structure prediction of biomolecular interactions with AlphaFold 3. Nature, 630:493–500.

Chai Discovery, Boitreaud J, Dent J, McPartlon M, Meier J, Reis V, Rogozhnikov A, Wu K. 2024. Chai-1: Decoding the molecular interactions of life. bioRxiv:2024.2010.2010.615955.

Herget S, Ranzinger R, Maass K, Lieth CW. 2008. GlycoCT-a unifying sequence format for carbohydrates. Carbohydr Res, 343:2162–2171.

Huang C, Kannan N, Moremen KW. 2025. Modeling glycans with AlphaFold 3: capabilities, caveats, and limitations. Glycobiology, 35.

Krishna R, Wang J, Ahern W, Sturmfels P, Venkatesh P, Kalvet I, Lee GR, Morey-Burrows FS, Anishchenko I, Humphreys IR, et al. 2024. Generalized biomolecular modeling and design with RoseTTAFold All-Atom. Science, 384:eadl2528.

Passaro S, Corso G, Wohlwend J, Reveiz M, Thaler S, Somnath VR, Getz N, Portnoi T, Roy J, Stark H, et al. 2025. Boltz-2: Towards Accurate and Efficient Binding Affinity Prediction. bioRxiv:2025.2006.2014.659707.

Stanley P, Moremen KW, Lewis NE, Taniguchi N, Aebi M. 2022. N-Glycans. In: Varki A, Cummings RD, Esko JD, Stanley P, Hart GW, Aebi M, Mohnen D, Kinoshita T, Packer NH, Prestegard JH, et al. editors. Essentials of Glycobiology. Cold Spring Harbor (NY): Cold Spring Harbor Laboratory Press. p. 103–116.

Tauriello G, Waterhouse AM, Haas J, Behringer D, Bienert S, Garello T, Schwede T. 2025. ModelArchive: A Deposition Database for Computational Macromolecular Structural Models. J Mol Biol, 437:168996.

Tiemeyer M, Aoki K, Paulson J, Cummings RD, York WS, Karlsson NG, Lisacek F, Packer NH, Campbell MP, Aoki NP, et al. 2017. GlyTouCan: an accessible glycan structure repository. Glycobiology, 27:915–919.

Tsuchiya S, Matsubara M, Aoki-Kinoshita KF, Yamada I. 2021. SugarDrawer: A Web-Based Database Search Tool with Editing Glycan Structures. Molecules, 26.

Varki A. 2017. Biological roles of glycans. Glycobiology, 27:3–49.

Varki A, Cummings RD, Aebi M, Packer NH, Seeberger PH, Esko JD, Stanley P, Hart G, Darvill A, Kinoshita T, et al. 2015. Symbol Nomenclature for Graphical Representations of Glycans. Glycobiology, 25:1323–1324.

Westbrook JD, Shao C, Feng Z, Zhuravleva M, Velankar S, Young J. 2015. The chemical component dictionary: complete descriptions of constituent molecules in experimentally determined 3D macromolecules in the Protein Data Bank. Bioinformatics, 31:1274–1278.

Woods RJ. 2018. Predicting the Structures of Glycans, Glycoproteins, and Their Complexes. Chem Rev, 118:8005–8024.

York WS, Mazumder R, Ranzinger R, Edwards N, Kahsay R, Aoki-Kinoshita KF, Campbell MP, Cummings RD, Feizi T, Martin M, et al. 2020. GlyGen: Computational and Informatics Resources for Glycoscience. Glycobiology, 30:72–73.

