## Supplementary Document 1 for "JAAG: a JSON input file Assembler for AlphaFold 3 with Glycan integration"

```
  JAAG HTML Input Structure Tree

  index.html
  ├── ABOUT JAAG SECTION
  │   └── Information & References Panel
  │
  ├── BASIC INFORMATION SECTION
  │   ├── Job Name
  │   └── Advanced Settings
  │        ├── JSON Input Version
  │        └── Model Seeds
  │             ├── Seed Count Generator
  │             └── Seeds List
  │
  ├── SEQUENCES SECTION
  │   │
  │   ├── Dynamic Sequence
  │   │   │
  │   │   ├── PROTEIN SEQUENCE INPUTS:
  │   │   │   ├── Chain ID
  │   │   │   ├── Copy Count
  │   │   │   ├── Protein Sequence
  │   │   │   │   ├── Manual Entry
  │   │   │   │   ├── UniProt ID
  │   │   │   │   └── Description
  │   │   │   ├── Glycosylation Options
  │   │   │   │   ├── Sequon Detection
  │   │   │   │   │   ├── Glycan Templates
  │   │   │   │   │   ├── Popup SugarDrawer for all Sequon
  │   │   │   │   │   └── Individual Sequon
  │   │   │   │   │        ├── Chain ID
  │   │   │   │   │        │   ├── Popup SugarDrawer
  │   │   │   │   │        │   └── GlycoCT String
  │   │   │   │   │        └── Description
  │   │   │   │   └── Manual Glycosylation Sites
  │   │   │   │        ├── Position
  │   │   │   │        ├── Residue
  │   │   │   │        ├── Linking Atom
  │   │   │   │        ├── Chain ID
  │   │   │   │        ├── Glycan Input
  │   │   │   │        │   ├── Popup SugarDrawer
  │   │   │   │        │   └── GlycoCT String
  │   │   │   │        └── Description
  │   │   │   ├── Modified Residues (PTM)
  │   │   │   │   ├── Position
  │   │   │   │   └── Modification (CCD)
  │   │   │   ├── User MSA
  │   │   │   │   ├── Automatic
  │   │   │   │   ├── File Paths
  │   │   │   │   │   ├── Unpaired MSA Path
  │   │   │   │   │   └── Paired MSA Path
  │   │   │   │   └── Direct Data
  │   │   │   │        ├── Unpaired MSA
  │   │   │   │        └── Paired MSA
  │   │   │   └── Structural Templates
  │   │   │        ├── Template Source Selection
  │   │   │        │   ├── mmCIF File Path
  │   │   │        │   │   └── File Path
  │   │   │        │   └── Inline mmCIF Data
  │   │   │        │        └── mmCIF Data
  │   │   │        ├── Query Indices
  │   │   │        └── Template Indices
  │   │   │
  │   │   ├── LIGAND SEQUENCE INPUTS:
  │   │   │   ├── Chain ID
  │   │   │   ├── Input Type
  │   │   │   │   ├── Glycan
  │   │   │   │   │   ├── Popup SugarDrawer
  │   │   │   │   │   └── GlycoCT String
  │   │   │   │   ├── CCD Codes
  │   │   │   │   │   └── CCD
  │   │   │   │   └── SMILES
  │   │   │   │        └── SMILES String
  │   │   │   ├── Copy Count
  │   │   │   └── Description
  │   │   │
  │   │   ├── RNA SEQUENCE INPUTS:
  │   │   │   ├── Chain ID
  │   │   │   ├── Copy Count
  │   │   │   ├── Sequence
  │   │   │   ├── Description
  │   │   │   ├── RNA Modifications
  │   │   │   │   ├── Position
  │   │   │   │   └── Modification (CCD)
  │   │   │   └── User MSA
  │   │   │        ├── Automatic
  │   │   │        ├── File Paths
  │   │   │        │   └── Unpaired MSA Path
  │   │   │        └── Direct Data
  │   │   │             └── Unpaired MSA
  │   │   │
  │   │   └── DNA SEQUENCE INPUTS:
  │   │        ├── Chain ID
  │   │        ├── Copy Count
  │   │        ├── Sequence
  │   │        ├── Description
  │   │        └── DNA Modifications
  │   │             ├── Position
  │   │             └── Modification (CCD)
  │   │
  │   └── bondedAtomPairs Section (pops up when adding ligand)
  │        ├── Chain 1 Dropdown
  │        ├── Residue 1 Dropdown
  │        ├── Atom 1
  │        ├── Chain 2 Dropdown
  │        ├── Residue 2 Dropdown
  │        └── Atom 2
  │
  ├── MANUAL USERCCD INPUT SECTION
  │   ├── Input Method Selection
  │   │   ├── Inline userCCD
  │   │   │   └── CIF
  │   │   └── userCCDPath
  │   │        └── CIF File Path
  │   ├── userCCD Code
  │   └── Active userCCDs List
  │
  └── JSON OUTPUT SECTION:
       ├── Lookup: Fetch GlyTouCan and GlyGen ID
       ├── Copy
       └── Download
```
